# Multi-omics analyses cannot identify true-positive novel associations from underpowered genome-wide association studies of four brain-related traits

**DOI:** 10.1101/2022.04.13.487655

**Authors:** David A.A. Baranger, Alexander S. Hatoum, Renato Polimanti, Joel Gelernter, Howard J. Edenberg, Ryan Bogdan, Arpana Agrawal

**Affiliations:** Department of Psychological & Brain Sciences, Washington University in St. Louis; Washington University School of Medicine, Department of Psychiatry, Saint Louis, USA; Department of Psychiatry, Division of Human Genetics, Yale School of Medicine, New Haven, CT, USA; Veterans Affairs Connecticut Healthcare System, West Haven, CT, USA; Department of Genetics, Yale School of Medicine, New Haven, CT, USA; Department of Neuroscience, Yale School of Medicine, New Haven, CT, USA; Department of Medical and Molecular Genetics, Indiana University School of Medicine, Indianapolis, IN, USA; Department of Biochemistry and Molecular Biology, Indiana University School of Medicine, Indianapolis, IN, USA

## Abstract

**Background:** The integration of multi-omics information (e.g., epigenetics and transcriptomics) can be useful for interpreting findings from genome-wide association studies (GWAS). It has additionally been suggested that multi-omics may aid in novel variant discovery, thus circumventing the need to increase GWAS sample sizes. We tested whether incorporating multi-omics information in earlier and smaller sized GWAS boosts true-positive discovery of genes that were later revealed by larger GWAS of the same/similar traits.

**Methods:** We applied ten different analytic approaches to integrating multi-omics data from twelve sources (e.g., Genotype-Tissue Expression project) to test whether earlier and smaller GWAS of 4 brain-related traits (i.e., alcohol use disorder/problematic alcohol use [AUD/PAU], major depression [MDD], schizophrenia [SCZ], and intracranial volume [ICV]) could detect genes that were revealed by a later and larger GWAS.

**Results:** Multi-omics data did not reliably identify novel genes in earlier less powered GWAS (PPV<0.2; 80% false-positive associations). Machine learning predictions marginally increased the number of identified novel genes, correctly identifying 1-8 additional genes, but only for well-powered early GWAS of highly heritable traits (i.e., ICV and SCZ). Multi-omics, particularly positional mapping (i.e., fastBAT, MAGMA, and H-MAGMA), was useful for prioritizing genes within genome-wide significant loci (PPVs = 0.5 – 1.0).

**Conclusions:** Although the integration of multi-omics information, particularly when multiple methods agree, helps prioritize GWAS findings and translate them into information about disease biology, it does not substantively increase novel gene discovery in brain-related GWAS. To increase power for discovery of novel genes and loci, increasing sample size is a requirement.

## Introduction

Genome-wide association studies (GWAS) have proven to be a uniquely effective tool for investigating the genetic architecture of complex traits. They have provided insights into the polygenic contributions of common variants, and yielded replicable genetic signals (1,2). However, for complex genetic traits, particularly for psychopathology, the number of cases needed for discovery is high, and depends on the heritability, polygenicity, heterogeneity, diagnostic accuracy, and prevalence of the trait (3,4). Schizophrenia, a highly heritable disorder, has witnessed linear increases in GWAS discoveries once a critical threshold was reached (5) (e.g., from 7 loci for 17,836 cases (6) to 270 loci for 69,369 cases (7)). Depression required a much larger sample size (5). Discoveries for substance use disorders have lagged, with evidence of greater polygenicity. Despite hundreds of loci for smoking behavior phenotypes (8), only 5 genome-wide significant loci have been identified for nicotine dependence (9). A similar broader non-diagnostic index of problematic alcohol use (2) yielded 29 loci, but findings are still quite limited for other drugs: only 10 loci for opioid use disorder (31,473 cases) (10) and 2 for cannabis use disorder (14,080 cases) (11). In parallel to GWAS, there has been an explosion in the multiomics data (e.g., gene expression, Hi-C) which are now available to interrogate GWAS findings. For some traits, expression data can also be drawn from model organisms in experimentally controlled exposure and behavioral paradigms (12). As greater evidence arises for genome-wide significant signals to be enriched in regulatory regions, these multi-omics data have proved to be valuable in gene prioritization. There has also been speculation that leveraging these ever-increasing omics sources might “recover” true signal from smaller GWAS, i.e., signals that may not meet criteria for genome-wide significance but are supported by multi-omics data, and eventually are identified as GWAS become larger, thus serving as a substitute for additional sample size (13–15).

We tested the hypothesis that the application of existing omics data and methods to a smaller-sized GWAS will yield additional “true positive” discoveries that would be found in the next, larger, GWAS. To test this hypothesis with respect to brain-related phenotypes, we selected four traits: (a) Alcohol use disorder/Problematic alcohol use (AUD/PAU), representing either diagnostic alcohol use disorder or an amalgam of diagnostic AUD and a non-diagnostic screener for problem drinking that is highly correlated with AUD; (b) Schizophrenia (SCZ), one of the earliest psychiatric disorders with many genome-wide significant findings that have increased with increasing sample size, making it an ideal reference trait for this test; (c) Major depression (MDD), a common, less heritable but highly polygenic and more clinically heterogeneous trait that is witnessing increasing discoveries in genome-wide significant loci, but at a much steeper cost of sample size. MDD is common, less heritable than SCZ and clinically heterogeneous and highly polygenic, similar to AUD/PAU; and (d) Intracranial volume (ICV), a highly heritable phenotype, which we selected to assess whether brain-derived omics data would be superior indices of genetic variability in the brain per se.

We addressed three primary hypotheses: (a) does incorporation of multiple omics (multiomics) sources to annotate the findings of the smaller GWAS recapitulate genes identified in the subsequent GWAS of the same or genetically closely-related trait (i.e., genetic correlation > 0.7), while minimizing false positives; (b) If multi-omics data can recover novel genes then which omics data and methods produce the most reliable predictions; and (c) does the multivariate consideration of these various sources of data improve prediction of genes? To test these, we examined sequential pairs of GWAS for a given trait (i.e., AUD/PAU, SCZ, MDD, ICV), where the latter GWAS, due to inclusion of more subjects, increased the number of genome-wide significant findings. Across analyses, we used genes, rather than variants, as the unit of discovery, because multi-omics data are gene-focused. We used a combination of omics data sources (including cross-species data) and statistical methods to annotate each GWAS. Key criteria were the positive predictive value (PPV, the proportion of genes identified in a larger GWAS relative to all genes predicted to be relevant using omics approaches in the prior, smaller GWAS), and the sensitivity (the proportion of genes identified by both the larger and smaller GWAS relative to all genes identified in the larger GWAS).

## Methods

### GWAS summary statistics

For each trait, analyses started with summary statistics from the smaller GWAS of the trait, ran a series of bioinformatic methods to identify additional genes, and compared the results to a subsequent GWAS of the trait with a larger sample size. Analyses focused on four traits: alcohol use disorder/problematic alcohol use (AUD/PAU) (1,2), major depressive disorder (MDD) (16,17), schizophrenia (SCZ) (7,18), and intracranial volume (ICV) (19). See **Table 1** for study details.

**Table 1.** Data sources. **A)** Publicly available summary statistics for each trait used in post-GWAS multi-omics analyses. * = Sample size omits data that is not publicly available. The number of significant risk loci (p<0.05×10^−8^) was calculated in FUMA, using publicly available summary statistics. **B)** Publicly available data used in post-GWAS multi-omics methods (see Table 2). **C)** Published data on genes which are differentially expressed in the brain, comparing participants with each disorder to controls. **D)** Published data on genes which are differentially expressed in mouse models of each disorder. **E)** Published rodent gene set on genes implicated in rodent models of alcohol use.

| **A: GWAS summary statistics** |  |  |  |
| --- | --- | --- | --- |
| Trait | Study | Sample size | # Genome-wide risk loci |
| Alcohol Use Disorder | Kranzler et al. 2019 (1) | N<sub>cases</sub>= 35,105<br>N<sub>control</sub>= 172,697 | 11 |
| Problematic Alcohol Use | Zhou et al. 2020 (2) | N<sub>effective</sub>= 300,789 | 29 |
| Major Depressive Disorder | Howard et al. 2019 (17) | N<sub>cases</sub>= 152,648*<br>N<sub>control</sub>= 562,035* | 51 |
| Major Depressive Disorder | Levey et al. 2021 (16) | N<sub>cases</sub>= 320,212*<br>N<sub>control</sub>= 581,929* | 100 |
| Schizophrenia | Ripke et al. 2014 (18) | N<sub>cases</sub>= 36,989<br>N<sub>control</sub>= 113,075 | 96 |
| Schizophrenia | Ripke et al. 2020 (7) | N<sub>cases</sub>= 69,369<br>N<sub>control</sub>= 236,642 | 242 |
| Intracranial Volume (UK Biobank only) | Jansen et al. 2020 (19) | N = 17,062 | 8 |
| Intracranial Volume | Jansen et al. 2020 (19) | N = 47,316 | 18 |
| **B: Post-GWAS multi-omics data sources** |  |  |  |
| Name/type | Study | Sample size | Notes |
| GTE<sub>x</sub> – brain RNA | GTE<sub>x</sub> Consortium, 2017 (31) | 80-154 | 13 brain regions |
| PsychEncode – brain RNA | Wang et al. 2018 (32) | 1,387 | 1 region |
| Brain-eMeta – brain RNA | Qi et al., 2018 (33) | 1,194 | Meta analysis across regions and samples |
| INTERVAL – plasma protein levels | Sun et al., 2018 (34) | 3,301 | - |
| **C: Human post-mortem brain gene expression** |  |  |  |
| Trait | Study | Case/Control | # Brain regions |
| Alcohol Use Disorder | Rao et al., 2020 (35) | 30/30 | 4 |
| Major Depressive Disorder | Wu et al., 2021 (36) | 101/96 | 7 |
| Schizophrenia | Collado-Torres et al., 2019 (37) | 286/265 | 2 |
| **D: Rodent post-mortem brain gene expression** |  |  |  |
| Trait | Study | # Models | # Brain regions |
| Alcohol Use Disorder | Ferguson et al., 2019 (38) | 1 | 7 |
| Major Depressive Disorder | Scarpa et al., 2020 (40) | 3 | 2 |
| Schizophrenia | Donegan et al., 2020 (55) | 3 | 1 |
| **E: Rodent gene set** |  |  |  |
| Trait | Study | Method | - |
| Alcohol Use Disorder | Huggett et al., 2021 (12) | GeneWeaver | - |

### Post-GWAS multi-omics methods

Eight post-GWAS multi-omics methods (**Table 2**) were used to identify genes associated with each trait, based on the results of each GWAS, including the max-SNP p-value (20), MAGMA (21), H-MAGMA (22), fastBAT (23), DEPICT (24), FUSION (25,26), S-MultiXcan (27), and SMR (28). These methods were selected as they are among the most widely-used and are representative of the major method types (i.e., positional: MAGMA, H-MAGMA, and fastBAT; expression-based: FUSION, S-MultiXcan, SMR, and DEPICT). These methods were applied to both data waves of each trait (i.e., both the larger and smaller GWAS). Analyses focused on gene-level associations, to incorporate information across methods and species. MAGMA was applied using the FUMA web platform (29), fastBAT, DEPICT, S-MultiXcan, and SMR were applied using the Complex-Traits Genetics Virtual Lab (30). Among the multi-omics methods used, FUSION and S-MultiXcan used RNA expression in the brain from the GTEx Consortium (31), and SMR used RNA expression in the brain from PsychENCODE(32) and the Brain-eMeta (33) study, as well as plasma protein expression from INTERVAL (34) (Table 1B).

**Table 2.** Post-GWAS multi-omics methods. Methods used to augment GWAS summary statistics and identify gene-level associations.

| Method | Description |
| --- | --- |
| SNP-based | Gene is assigned the p-value of the most significant SNP within 10kb of its boundaries (20). |
| MAGMA | Set-based analysis, regression-based p-values (21). |
| H-MAGMA | Incorporates chromatin interaction profiles for gene boundaries (22) . |
| fastBAT | Fast set-based analysis, simulation-based p-values (23). |
| DEPICT | Predicts gene function to prioritize the most likely causal genes (24). |
| FUSION | Transcriptome-wide association study (TWAS) using GTEx gene expression to impute gene expression. Integrates across multiple tissues using an aggregated Cauchy association test (25,26). |
| S-MultiXcan | Transcriptome-wide association study (TWAS) using GTEx gene expression to impute gene expression. Integrates across multiple tissues using multivariate regression (27). |
| SMR | Summary mendelian randomization. Tests for mediation of genetic effects by gene or protein expression, using Brain-eMeta, PsychEncode, and INTERVAL (28). |

### Additional multi-omics data

Additional multi-omics data were drawn from published data sets: genes differentially expressed in human brain tissue (Table 1C) (35–37) and genes differentially expressed in brains of mouse models of each disorder (Table 1D). For AUD/PAU, data came from alcohol-naïve mice from a line bred for binge-drinking-like behavior, High Drinking in the Dark mice (38). Results from 7 tissues were integrated using an aggregated Cauchy association test (39), which is also used by the FUSION TWAS method to integrate across tissues (25). An additional list of genes from rodent studies enriched for the heritability of alcohol use disorder was included as an additional method (12). MDD-associated genes came from a study examining differentially expressed genes in two brain tissues across three mouse chronic stress models (40). Results within each model were similarly combined across tissues using an aggregated Cauchy association test. As results were only reported for nominally significant genes, precluding a meta-analysis combining models, the minimum p-value across the three models was assigned as a gene-level p-value. SCZ-associated genes came from a study examining differentially expressed genes in one brain tissue across three mouse models of developmental disruption. Results were combined across models using Fisher’s combined probability test (41).

### Predictive value of significant genes

Multi-omics methods were applied to summary data from the smaller GWAS. Genes identified by each multi-omics method were defined as those surviving false discovery rate correction within method. Genes that were either not measured in a data source (e.g., RNA expression was too low to measure), or unreported (i.e., some data sources only reported genes that were at least nominally-significant) were marked as showing no association for that source. For each trait, the target set of genes was defined as genes close (i.e., ±10 kb) to a genome-wide significant locus (p<5×10^−8^) in the larger of the two paired GWAS.

Analyses for novel gene discovery examined genes that were *not* proximal to a genome-wide significant locus in the smaller GWAS but *were* identified by individual multi-omics methods or combinations of methods. These analyses tested whether these genes were identified as genome-wide significant in the *larger* GWAS. Analyses for prioritization examined genes identified by proximity to a genome-wide significant locus in the *smaller* GWAS. These analyses tested if these genes were more likely to be identified as genome-wide significant in the *larger* GWAS if they were also identified by individual multi-omics methods or combinations of methods.

Two primary performance metrics were used, the positive predictive value (PPV) and sensitivity (42). The PPV, which ranges from 0-1, reflects the probability that a positive prediction reflects a true genetic signal (i.e., the ratio of the true positive rate to the sum of the true and false positive rates). PPV reflects what proportion of target genes identified in the later, larger, GWAS, are genes predicted to be relevant using omics approaches in the prior, smaller GWAS. For example, a PPV of 0.75 would mean that 75% of the genes identified by multi-omics data in the smaller GWAS are positionally significant in the larger GWAS. Sensitivity, which also ranges from 0-1, reflects the proportion of the positionally significant genes in the larger GWAS captured by the test (the ratio of the true positive rate to the sum of the true positive and false negative rates). That is, of the genes positionally identified in the later, larger, GWAS, what proportion are found using multi-omics data in the prior, smaller, GWAS? For example, a sensitivity of 0.5 would mean that multi-omics identified 50% of all the positionally significant genes in the later GWAS. Thus, an ideal test would have both a high PPV and a high sensitivity (i.e., the test captures the majority of the significant genes in the larger GWAS, with very few incorrect predictions). A test with a high PPV and low sensitivity misses the majority of the significant genes in the larger GWAS, but the few that are identified are mostly correct predictions. Conversely, a test with high sensitivity and a low PPV captures the majority of the significant genes in the larger GWAS, but at the cost of a high number of incorrect predictions.

### Multivariate machine learning gene prioritization

#### Training procedure

As the predictions of individual methods are not perfectly correlated (**Supplemental Figure 1**), and methods may only partially contribute to prediction (i.e., may need to be weighted), a multivariate combination of methods might improve performance. Models were trained, using only the earlier GWAS, to predict which genes were positionally linked to genome-wide significant SNPs using data from multi-omics methods. Summary data in the smaller GWAS were split into a training and testing component. We estimated PPV and sensitivity using the hold-out (testing) set, and then subsequently in the larger future GWAS. For example, in MDD, ML models were trained using half the chromosomes from Howard et al. (17). These models were then tested in the remaining chromosomes of Howard et al., which yielded a predicted probability of how likely a gene is to be proximal to a genome-wide significant locus, based on multi-omics data. A probability threshold was selected by examining PPV and sensitivity distributions in Howard et al. We then took the genes which surpassed the identified threshold and assessed the PPV and sensitivity of those predictions in the larger Levey et al. (16) GWAS.

#### ML Algorithms

GWAS of psychiatric disorders and brain-related traits are currently only powered to find small proportions of variants associated with these traits. Standard ML classifiers may thus not be appropriate, as these methods perform best when data are evenly balanced (i.e., when 50% of genes are significant). Therefore, we used a combination of up-sampling, bootstrapping, bagging, and ensemble learning in two algorithms. Missing data across methods was handled with a median impute (bagimpute and K-means impute were unreliable due to high missingness). The first method was AdaBoosting (43); a tree ensemble method that was originally developed to predict rare outcomes (44). AdaBoosted trees were bagged with up-sampling minority cases. The second method was model-average neural network (ANN) (45); a series of simple functions which attempt to capture patterns in the data. The model ANN averages across many runs to develop a prediction. An ensemble prediction was then generated by averaging the predictions from the two approaches. All machine learning models were trained with 3-fold cross-validation with up-sampling of genes containing GWAS-significant SNPs. This cross-validation and training was repeated 10 times (bootstrapping with replacement), increasing the proportion of genes containing GWAS-significant SNPs in the training data.

Finally, we determined feature importance to identify which multi-omics methods most contributed to gene identification. For ANN, an ROC curve was generated for each variable using sensitivity and specificity, and the probabilities were cut off at a series of (random) points. Using the trapezoidal rule, the area under the ROC was calculated for each multi-omics method and used as a measure of feature importance (46). When using AdaBoost we used tree-specific feature importance, computed by summing how much the model improved each time it used a multi-omics method (47).

## Results

Correlations between gene sets identified by multi-omics were quite low (median=0.12, range=0-0.7; **Supplementary Figure 1**). Related methods identified more similar gene sets, though correlations remained moderate (positional: median=0.4, range=-.27-0.63; expression-based: median=0.38, range=0.22-0.72). Genes identified by methods using GWAS summary statistics showed very low overlap with genes identified by post-mortem gene expression studies in humans and rodents and with rodent-based gene set analyses (range=-0.01-0.02).

For novel gene discovery (**Figure 1**), no method achieved a PPV greater than 0.2 for any trait (i.e., 80% of genes identified by multi-omics were *not* proximal to a genome-wide significant locus in the larger GWAS) with the exception of fastBAT for AUD/PAU, which correctly identified a single gene (*ADH1A*), yielding a PPV of 1.0 (**Figure 1A**). Similarly, upon examination of agreement across methods, none surpassed a PPV of 0.22 for any trait (**Figure 1B**).

**Figure 1.**
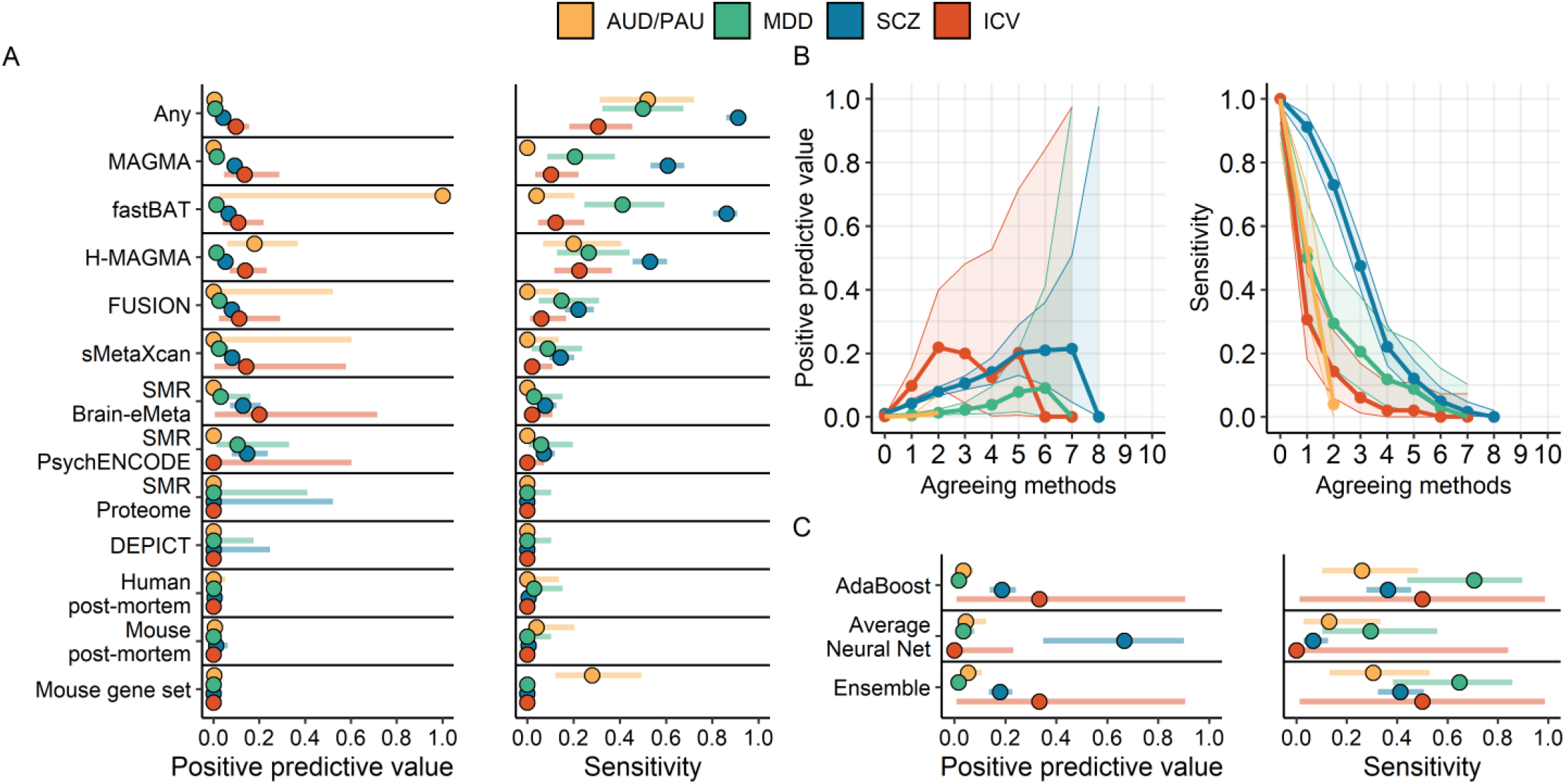
Multi-omics cannot reliably identify true-positive novel genes. Genes that are not proximal to a GWAS-significant locus but are identified by multi-omics methods in the smaller GWAS are not likely to contain a GWAS-significant locus in the larger GWAS. A) The positive predictive value and sensitivity for each method, and for all methods combined (Any). B) Performance for increasing agreement between multi-omics methods. C) Performance of machine-learning (ML) in the larger GWAS (ML was trained in the smaller GWAS). Points represent the estimates, while horizonal bars reflect the 95% CI. AUD/PAU = Alcohol use disorder/Problematic alcohol use; MDD = Major depressive disorder; SCZ = Schizophrenia; ICV = Intracranial volume.

For gene prioritization, genes that were identified both by proximity to a genome-wide significant locus and by multi-omics were more likely to be significant in the larger GWAS than genes only identified by a genome-wide significant locus (**Figure 2A**). However, no method had both a high PPV and a high sensitivity for any trait. Expression-based methods had a slightly higher PPV than position-based methods, but position-based methods attained a much greater sensitivity for all traits. Analyses examining agreement between methods found that genes identified by more methods (regardless of the specific method) had a higher PPV across all traits, though with a lower sensitivity (**Figure 2B**). That is, when more methods agreed, the gene was more likely to be positionally significant in the larger GWAS. However, the low sensitivity indicates that this approach misses the majority of findings in the larger GWAS.

**Figure 2.**
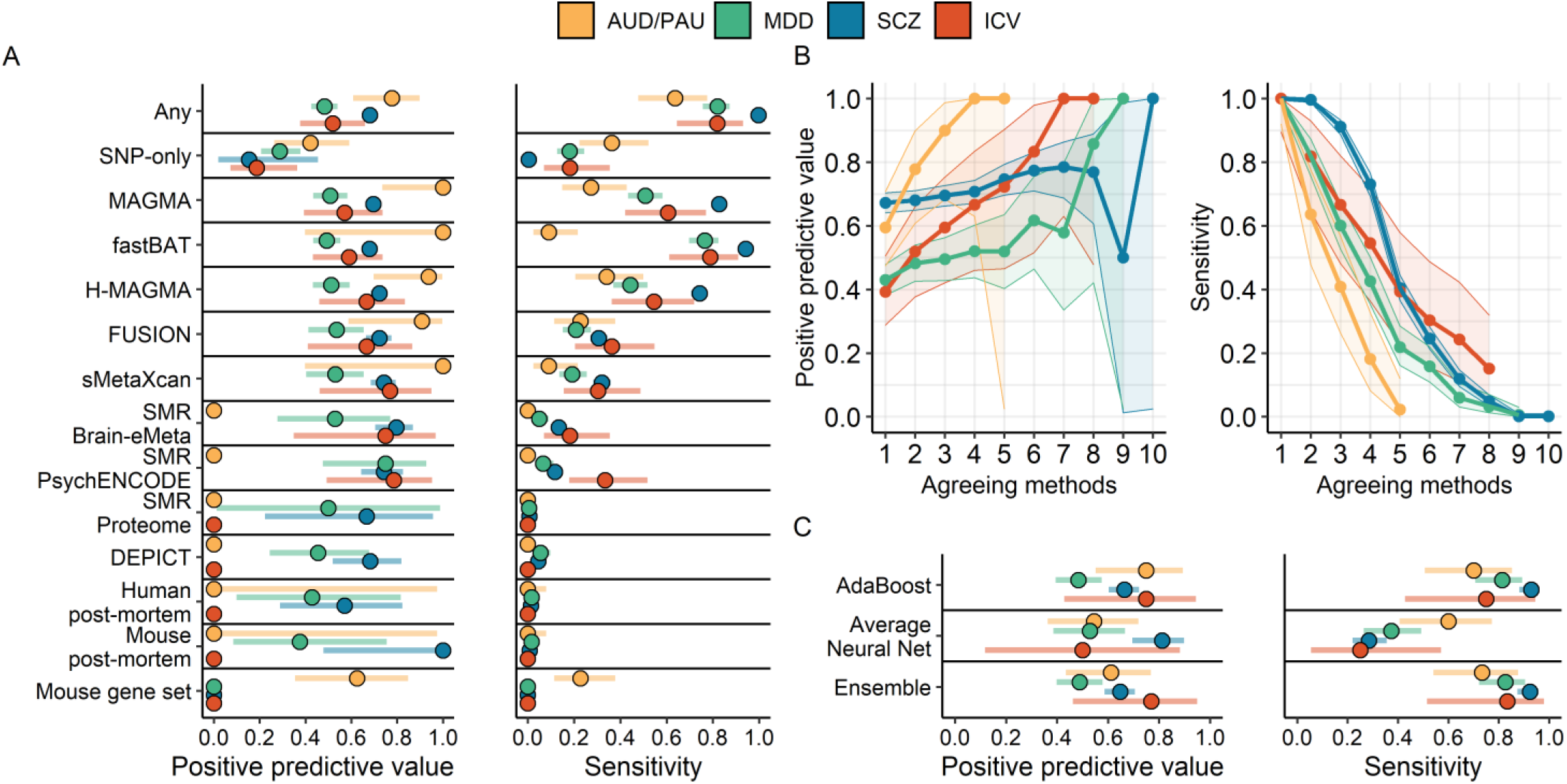
Multi-omics is useful for prioritization of GWAS-significant genes. Genes that are proximal to a GWAS-significant locus and are identified by multi-omics methods in the smaller GWAS are likely to contain a GWAS-significant locus in the larger GWAS. A) The positive predictive value and sensitivity for each method, and for all methods combined (Any). ‘SNP-only’ = the gene is proximal to a GWAS-significant locus, but is not identified by multi-omics. B) Performance for increasing agreement between multi-omics methods. C) Performance of machine-learning (ML) in the larger GWAS (ML was trained in the smaller GWAS). Points represent the estimates, while horizonal bars reflect the 95% CI. AUD/PAU = Alcohol use disorder/Problematic alcohol use; MDD = Major depressive disorder; SCZ = Schizophrenia; ICV = Intracranial volume.

Across both novel gene discovery and gene prioritization analyses, genes identified by more methods in the smaller GWAS tended to have a higher -log10(p) value in the larger GWAS and similarly tended to be identified by more methods in the larger GWAS (Supplemental Results; **Supplemental Figures 2&3**). Supplemental analyses additionally examined performance across all possible combinations of multi-omics methods, as well as for different definitions of a ‘significant’ gene in the larger GWAS. No combination performed best across all traits, and the pattern of associations for different definitions of ‘significant’ (i.e., p<5×10^−6^ or identified by multi-omics in the larger GWAS) remained similar (Supplemental Results; **Supplemental Figures 4&5**).

### Machine learning

Model predictions are probability scores (i.e., the probability that a gene contains a genome-wide significant SNP, based on multi-omics). The performance (PPV and sensitivity) of different cut-offs was evaluated in the hold-out data from the smaller GWAS (**Supplemental Figure 6**). A probability cut-off of 75% was selected for AdaBoost, 90% for the model-average neural net, and 50% for the ensemble of the two. The performance of each gene set was then assessed in the later, larger GWAS (**Supplemental Table 1**). ML performance in the held-out data from the smaller GWAS was moderately reflective of performance in the larger GWAS (**Supplemental Figure 7 A&B**), with the exception of AUD/PAU. Novel gene discovery (**Figure 1C**) was the most successful for SCZ (PPV = 0.67), though only a minority of novel genes were found (sensitivity = 0.066; i.e., 8 genes). Similarly, for ICV, one of the three identified genes was significant. Novel gene discovery in AUD/PAU and MDD did not surpass a PPV of 0.1. AdaBoost attained moderate performance for gene prioritization across all traits (PPV = 0.48-0.75, sensitivity = 0.42–0.88; **Figure 2C**). While the average neural net had a higher PPV for SCZ (0.8), this was at the cost of much lower sensitivity (0.28). Examination of variable importance scores revealed that fastBAT, MAGMA, and H-MAGMA were the primary contributors to prediction across models and methods (**Figure 3**). DEPICT also attained a comparable level of importance for AUD/PAU in the average neural net and for MDD in AdaBoost. In general, gene expression methods were the least informative predictors.

**Figure 3.**
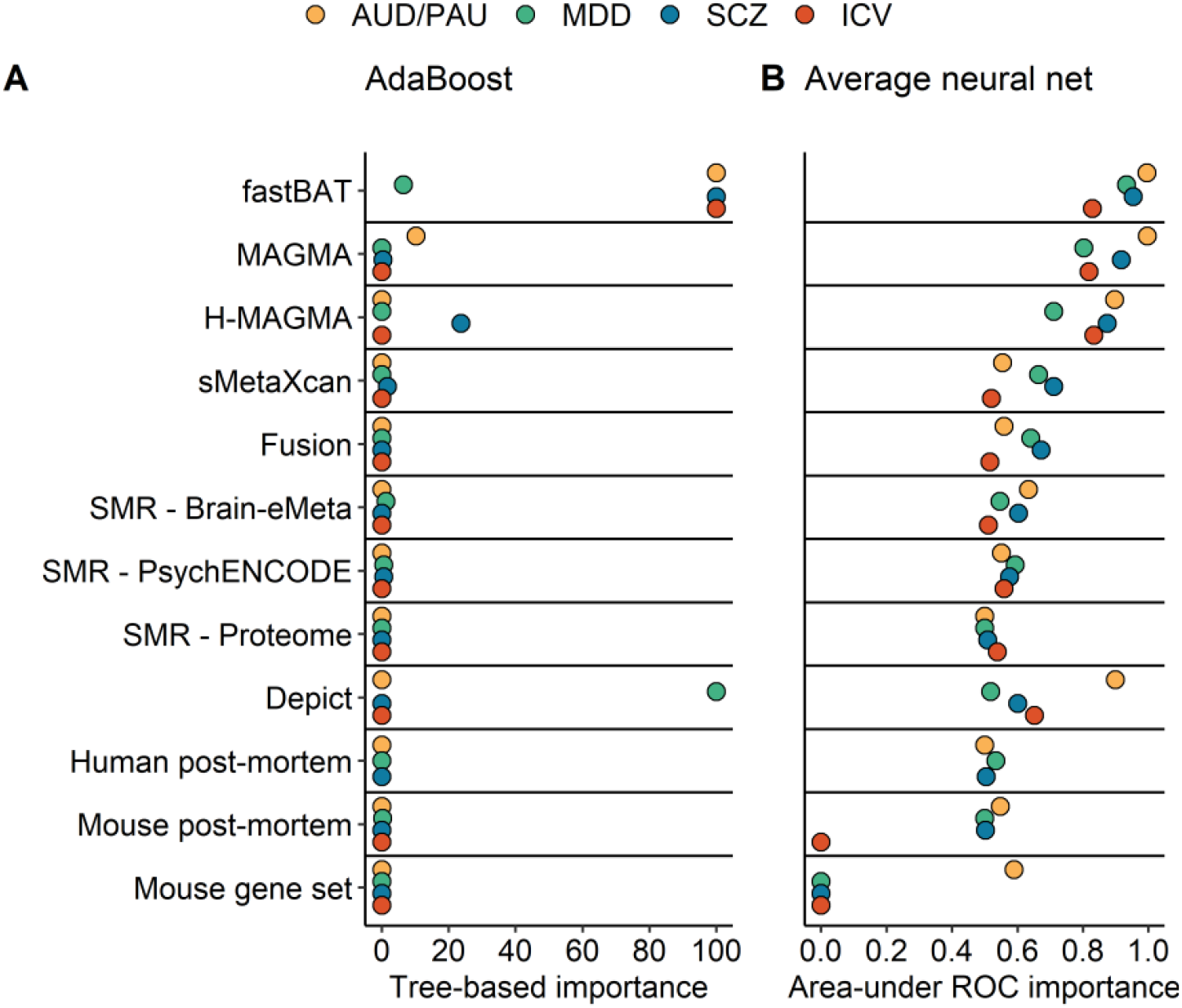
Machine learning variable importance. A) Average neural net variable importance using the area-under the ROC method. The maximum possible importance is ‘1.0’. B) Tree-based importance in AdaBoost. The maximum possible importance is ‘100’. AUD/PAU = Alcohol use disorder/Problematic alcohol use; MDD = Major depressive disorder; SCZ = Schizophrenia; ICV = Intracranial volume.

## Discussion

GWAS have provided unique insight into the genetic architecture of complex traits. Their informativeness, particularly for complex and heterogeneous traits, is tightly dependent on sample size. As increasing GWAS sample size is arduous and expensive, there is a natural desire to identify additional sources of data that may increase signal within existing GWAS data. In particular, it has been hypothesized that augmenting GWAS with other kinds of *omics* data (e.g., transcriptomics and data from non-human animal models) may boost signal within existing less powered GWAS to circumvent the need for additional GWAS data. In contrast to this hypothesis, we found that current omics approaches, in the context of existing GWAS sample sizes, does not foreshadow later discoveries by larger GWAS. Increasing GWAS sample size is a requirement to increase power for discovery of novel genes and loci.

Using a smaller GWAS and a subsequent larger GWAS of the same trait, we examined whether applying multi-omics to gene identification in the smaller GWAS could identify genes that will become significant in a future larger GWAS. No single method or combination of methods achieved both a high PPV and sensitivity when “predicting” novel signals in a larger subsequent GWAS (**Figure 1**). While machine learning out-performed individual methods and even linear combinations of methods, it only correctly identified an additional 1-8 genes, and only for highly heritable traits with a well-powered smaller GWAS (SCZ and ICV). This observation converges with related work evaluating methods for SNP and locus annotation (13,14,48), which has concluded that such methods can only marginally increase the number of true-positive observations. Similarly, prior work examining positional gene-based methods in a simulated trait with relatively few causal SNPs (n=602) observed a tradeoff between sensitivity and specificity (49), which is also seen here. Overall, multi-omics data for novel gene discovery incur either a high false-negative (i.e., they miss many novel discoveries) or a high false-positive burden (i.e., they have many findings which are not present when the sample size increases), and are thus not a reasonable replacement for larger discovery samples.

As expected, GWAS-significant genes that are also prioritized by multi-omics were more likely to replicate in future GWAS (**Figure 2**). However, the number of agreeing methods necessary to attain even a moderate PPV (i.e., PPV = 0.6) differed for each trait (1-6), and no single method or combination of methods performed the best across all traits (**Figure 2, Supplemental Figure 4**). Machine learning (ML) performed similarly to the best-performing individual and combinations of methods. Broadly, these results support current practices in the field, wherein associations that are GWAS-significant and robust across multi-omics methods are given the greatest credence, and suggest that further development of ML approaches to post-GWAS multi-omics integration may be a fruitful avenue for prioritization within loci (50).

Despite the variation in method performance, positional methods (i.e., fastBAT, MAGMA, and H-MAGMA) were frequently among the top performers. These achieved a comparable PPV to expression-based methods, with superior sensitivity. Positional methods were also the top ranked by both machine learning algorithms (with the exception of MDD analyses with AdaBoost, where fastBAT was second to DEPICT; Figure 3). Analyses defined an association in the larger GWAS as those that are proximal (within 10 kb) to a GWAS-significant SNP, which could have biased results towards position-based methods. This definition was selected as the most-proximal gene has been found to be the most accurate method in cases where associated genes are known and established (51). However, it is an imperfect approach, as many significant SNPs are not proximal to a protein-coding region (51,52). Indeed, this observation was a major impetus for the development of the expression-based multi-omics methods used here (22,26). Follow-up analyses thus tested whether multi-omics could identify genes that would be Bonferroni-significant in a similar multi-omics analysis of the larger GWAS (**Supplemental Figure 8**). Expression-based methods largely achieved a higher PPV than the position-based methods (though not for AUD/PAU), but at the cost of much lower sensitivity. This result suggests that expression-based methods may have more method-specific variance in their results, leading to enhanced within-method prediction and reduced cross-method prediction. This again emphasizes our observation that agreement between methods leads to more reliable results.

Across analyses, the success of multi-omics approaches appeared largely to depend on trait heritability and GWAS power. Analyses were the most successful in SCZ (the best-powered GWAS), attaining the largest PPV and sensitivity. However, despite a relatively large sample size, analyses were largely the least successful in MDD (the trait with the lowest SNP-based heritability). AUD/PAU was an occasional exception to this general pattern, wherein PPV greatly increased for each additional agreeing method for gene prioritization (**Figure 2**). Substance use disorders are unique among psychiatric disorders, as there are a few large-effect loci mapping to substance-specific receptors and bioavailability pathways (53). The increase in AUD/PAU PPV likely reflects the relatively large effect size of genes in these pathways, and indeed it largely included genes implicated in alcohol metabolism (i.e., *ADH1B, ADH5, and ADH7*). However, these results more broadly suggest that multi-omics will be the least able to successfully prioritize findings, and to identify novel associations, in traits where it would be the most useful (i.e., where GWAS are particularly underpowered and no loci of large effect are evident).

Analyses focused on trait associations at the gene level, rather than individual SNPs (14). This enabled the additional integration of multi-omics data that did not use GWAS information, including results from post mortem studies of gene expression in patients and rodent models, and a gene-set for AUD/PAU that integrates information from a variety of rodent data sources (12). These methods were uniformly the least informative across all analyses. However, data from post-mortem studies of MDD and SCZ, and the AUD/PAU rodent gene set, had a moderate PPV and low sensitivity for gene prioritization. We note that we did not systematically query the literature to derive comprehensive gene-sets for all traits. Indeed, the AUD/PAU rodent gene set, which was derived from a systematic review of the literature (12), achieved a higher PPV and sensitivity for AUD/PAU than data from individual studies of rodent models did for their respective traits. The gene-level focus reflects an additional weakness of the omics-integration approach, in that it could lead to improved knowledge of biology without narrowing down the identity of causal loci in human populations. Thus, while recent related studies using locus-level analyses yielded similar findings (14), we cannot rule out that alternative methods could have led to stronger results.

We note some additional limitations of the present study. Analyses only used GWAS from European samples. Recent work suggests that gene-level findings from expression-based methods may be more replicable across ancestries than SNP-level effects (54). Thus, expression-based methods may have superior performance in contexts that could not be evaluated in the present study, owing to the lack of well-powered GWAS for brain-related traits in non-European samples. We tested four different traits that are quite different, to cover a range of trait characteristics, but we cannot exclude that some other traits might respond more or less successfully to the integration of omics data. Lastly, the smaller GWAS were subsets of the subsequent larger GWAS. This should have favorably biased the tests, but nevertheless they did not perform well.

## Conclusions

The present results demonstrate that multi-omics are not a replacement for increasing GWAS sample size. Instead, results support the use of multi-omics as methods for prioritizing genes that contain GWAS-significant loci. Underpowered traits (e.g., cannabis use disorder (11)) will require much larger sample sizes before even prioritization will be possible. We view the combination of larger GWAS sample sizes and multi-omics method advancements as likely the most fruitful avenue for identifying multiple plausible causal loci for brain-related traits.

## References

1. Kranzler HR, Zhou H, Kember RL, Vickers Smith R, Justice AC, Damrauer S, et al. (2019): Genome-wide association study of alcohol consumption and use disorder in 274,424 individuals from multiple populations. Nat Commun 10: 1499.

2. Zhou H, Sealock JM, Sanchez-Roige S, Clarke T-K, Levey DF, Cheng Z, et al. (2020): Genome-wide meta-analysis of problematic alcohol use in 435,563 individuals yields insights into biology and relationships with other traits. Nat Neurosci 23: 809–818.

3. Cai N, Revez JA, Adams MJ, Andlauer TFM, Breen G, Byrne EM, et al. (2020): Minimal phenotyping yields genome-wide association signals of low specificity for major depression [no. 4]. Nat Genet 52: 437–447.

4. Visscher PM, Wray NR, Zhang Q, Sklar P, McCarthy MI, Brown MA, Yang J (2017): 10 Years of GWAS Discovery: Biology, Function, and Translation. Am J Hum Genet 101: 5–22.

5. Sullivan PF, Geschwind DH (2019): Defining the Genetic, Genomic, Cellular, and Diagnostic Architectures of Psychiatric Disorders. Cell 177: 162–183.

6. Schizophrenia Psychiatric Genome-Wide Association Study (GWAS) Consortium. Genome-wide association study identifies five new schizophrenia loci. Nat Genet. 2011 Sep 18;43(10):969–76.

7. The Schizophrenia Working Group of the Psychiatric Genomics Consortium, Ripke S, Walters JTR, O’Donovan MC, et al. (2022): Mapping genomic loci prioritises genes and implicates synaptic biology in schizophrenia. medRxiv 2020.09.12.20192922.

8. Liu M, Jiang Y, Wedow R, Li Y, Brazel DM, Chen F, et al. (2019): Association studies of up to 1.2 million individuals yield new insights into the genetic etiology of tobacco and alcohol use. Nat Genet 51: 237–244.

9. Quach BC, Bray MJ, Gaddis NC, Liu M, Palviainen T, Minica CC, et al. (2020): Expanding the genetic architecture of nicotine dependence and its shared genetics with multiple traits [no. 1]. Nat Commun 11: 5562.

10. Kember RL, Vickers-Smith R, Xu H, Toikumo S, Niarchou M, Zhou H, et al. (2021, December 15): Cross-ancestry meta-analysis of opioid use disorder uncovers novel loci with predominant effects on brain. medRxiv, p 2021.12.13.21267480.

11. Johnson EC, Demontis D, Thorgeirsson TE, Walters RK, Polimanti R, Hatoum AS, et al. (2020): A large-scale genome-wide association study meta-analysis of cannabis use disorder. Lancet Psychiatry 7: 1032–1045.

12. Huggett SB, Johnson EC, Hatoum AS, Lai D, Bubier JA, Chesler EJ, et al. (2021): Genes Identified in Rodent Studies of Alcohol Intake Are Enriched for Heritability of Human Substance Use. p 2021.03.22.436527.

13. Pickrell JK (2014): Joint analysis of functional genomic data and genome-wide association studies of 18 human traits. Am J Hum Genet 94: 559–573.

14. Moore A, Marks J, Quach BC, Guo Y, Bierut LJ, Gaddis NC, et al. (2022, January 11): Evaluation of methods incorporating biological function and GWAS summary statistics to accelerate discovery. bioRxiv, p 2022.01.10.475153.

15. Ayalew M, Le-Niculescu H, Levey DF, Jain N, Changala B, Patel SD, et al. (2012): Convergent functional genomics of schizophrenia: from comprehensive understanding to genetic risk prediction [no. 9]. Mol Psychiatry 17: 887–905.

16. Levey DF, Stein MB, Wendt FR, Pathak GA, Zhou H, Aslan M, et al. (2021): Bi-ancestral depression GWAS in the Million Veteran Program and meta-analysis in >1.2 million individuals highlight new therapeutic directions. Nat Neurosci 24: 954–963.

17. Howard DM, Adams MJ, Clarke T-K, Hafferty JD, Gibson J, Shirali M, et al. (2019): Genome-wide meta-analysis of depression identifies 102 independent variants and highlights the importance of the prefrontal brain regions. Nat Neurosci 22: 343–352.

18. Ripke S, Neale BM, Corvin A, Walters JTR, Farh K-H, Holmans P a., et al. (2014): Biological insights from 108 schizophrenia-associated genetic loci. Nature 511: 421–427.

19. Jansen PR, Nagel M, Watanabe K, Wei Y, Savage JE, de Leeuw CA, et al. (2020): Genome-wide meta-analysis of brain volume identifies genomic loci and genes shared with intelligence. Nat Commun 11: 5606.

20. Helgeland Ø (2019): Oyhel/Vautils. Retrieved December 15, 2021, from https://github.com/oyhel/vautils

21. de Leeuw CA, Mooij JM, Heskes T, Posthuma D (2015): MAGMA: Generalized Gene-Set Analysis of GWAS Data. PLoS Comput Biol 11: 1–19.

22. Sey NYA, Hu B, Mah W, Fauni H, McAfee JC, Rajarajan P, et al. (2020): A computational tool (H-MAGMA) for improved prediction of brain-disorder risk genes by incorporating brain chromatin interaction profiles. Nat Neurosci 23: 583–593.

23. Bakshi A, Zhu Z, Vinkhuyzen AAE, Hill WD, McRae AF, Visscher PM, Yang J (2016): Fast set-based association analysis using summary data from GWAS identifies novel gene loci for human complex traits. Sci Rep 6: 32894.

24. Pers TH, Karjalainen JM, Chan Y, Westra H-J, Wood AR, Yang J, et al. (2015): Biological interpretation of genome-wide association studies using predicted gene functions. Nat Commun 6: 5890.

25. Feng H, Mancuso N, Gusev A, Majumdar A, Major M, Pasaniuc B, Kraft P (2021): Leveraging expression from multiple tissues using sparse canonical correlation analysis and aggregate tests improves the power of transcriptome-wide association studies. PLOS Genet 17: e1008973.

26. Gusev A, Ko A, Shi H, Bhatia G, Chung W, Penninx BWJH, et al. (2016): Integrative approaches for large-scale transcriptome-wide association studies. Nat Genet 48: 245–252.

27. Barbeira AN, Pividori M, Zheng J, Wheeler HE, Nicolae DL, Im HK (2019): Integrating predicted transcriptome from multiple tissues improves association detection. PLOS Genet 15: e1007889.

28. Zhu Z, Zhang F, Hu H, Bakshi A, Robinson MR, Powell JE, et al. (2016): Integration of summary data from GWAS and eQTL studies predicts complex trait gene targets. Nat Genet 48: 481–487.

29. Watanabe K, Taskesen E, van Bochoven A, Posthuma D (2017): Functional mapping and annotation of genetic associations with FUMA. Nat Commun 8: 1826.

30. Cuéllar-Partida G, Lundberg M, Kho PF, D’Urso S, Gutiérrez-Mondragón LF, Ngo TT, Hwang L-D (2019): Complex-Traits Genetics Virtual Lab: A Community-Driven Web Platform for Post-GWAS Analyses. p 518027.

31. Aguet F, Brown AA, Castel SE, Davis JR, He Y, Jo B, et al. (2017): Genetic effects on gene expression across human tissues. Nature 550: 204–213.

32. Wang D, Liu S, Warrell J, Won H, Shi X, Navarro FCP, et al. (2018): Comprehensive functional genomic resource and integrative model for the human brain. Science 362: eaat8464.

33. Qi T, Wu Y, Zeng J, Zhang F, Xue A, Jiang L, et al. (2018): Identifying gene targets for brain-related traits using transcriptomic and methylomic data from blood. Nat Commun 9: 2282.

34. Sun BB, Maranville JC, Peters JE, Stacey D, Staley JR, Blackshaw J, et al. (2018): Genomic atlas of the human plasma proteome. Nature 558: 73–79.

35. Rao X, Thapa KS, Chen AB, Lin H, Gao H, Reiter JL, et al. (2021): Allele-specific expression and high-throughput reporter assay reveal functional genetic variants associated with alcohol use disorders. Mol Psychiatry 26: 1142–1151.

36. Wu W, Howard D, Sibille E, French L (2021): Differential and spatial expression meta-analysis of genes identified in genome-wide association studies of depression. Transl Psychiatry 11: 1–12.

37. Collado-Torres L, Burke EE, Peterson A, Shin J, Straub RE, Rajpurohit A, et al. (2019): Regional Heterogeneity in Gene Expression, Regulation, and Coherence in the Frontal Cortex and Hippocampus across Development and Schizophrenia. Neuron 103: 203–216.e8.

38. Ferguson LB, Zhang L, Kircher D, Wang S, Mayfield RD, Crabbe JC, et al. (2019): Dissecting Brain Networks Underlying Alcohol Binge Drinking Using a Systems Genomics Approach. Mol Neurobiol 56: 2791–2810.

39. Liu Y, Chen S, Li Z, Morrison AC, Boerwinkle E, Lin X (2019): ACAT: A Fast and Powerful p Value Combination Method for Rare-Variant Analysis in Sequencing Studies. Am J Hum Genet 104: 410–421.

40. Scarpa JR, Fatma M, Loh Y-HE, Traore SR, Stefan T, Chen TH, et al. (2020): Shared Transcriptional Signatures in Major Depressive Disorder and Mouse Chronic Stress Models. Biol Psychiatry 88: 159–168.

41. Moulos P, Hatzis P (2015): Systematic integration of RNA-Seq statistical algorithms for accurate detection of differential gene expression patterns. Nucleic Acids Res 43: e25.

42. Stevenson M, Sergeant E, Nunes T, Heuer C, Marshall J, Sanchez J, et al. (2021): EpiR: Tools for the Analysis of Epidemiological Data, version 2.0.40. Retrieved December 16, 2021, from https://CRAN.R-project.org/package=epiR

43. Alfaro E, Gamez M, García N (2013): adabag: An R Package for Classification with Boosting and Bagging. J Stat Softw 54: 1–35.

44. Rojas R (2009): AdaBoost and the super bowl of classifiers a tutorial introduction to adaptive boosting. Freie Univ Berl Tech Rep.

45. Venables WN, Ripley BD (2002): Modern Applied Statistics with S ((W. N. Venables & B. D. Ripley, editors)). New York, NY: Springer. https://doi.org/10.1007/978-0-387-21706-2_1

46. Kuhn M (2008): Building Predictive Models in R Using the caret Package. J Stat Softw 28: 1–26.

47. Friedman JH (2001): Greedy function approximation: A gradient boosting machine. Ann Stat 29: 1189–1232.

48. Dey KK, van de Geijn B, Kim SS, Hormozdiari F, Kelley DR, Price AL (2020): Evaluating the informativeness of deep learning annotations for human complex diseases. Nat Commun 11: 4703.

49. Wojcik GL, Kao WL, Duggal P (2015): Relative performance of gene-and pathway-level methods as secondary analyses for genome-wide association studies. BMC Genet 16: 34.

50. Nicholls HL, John CR, Watson DS, Munroe PB, Barnes MR, Cabrera CP (2020): Reaching the End-Game for GWAS: Machine Learning Approaches for the Prioritization of Complex Disease Loci. Front Genet 11. Retrieved February 22, 2022, from https://www.frontiersin.org/article/10.3389/fgene.2020.00350

51. Nasser J, Bergman DT, Fulco CP, Guckelberger P, Doughty BR, Patwardhan TA, et al. (2021): Genome-wide enhancer maps link risk variants to disease genes [no. 7858]. Nature 593: 238–243.

52. Maurano MT, Humbert R, Rynes E, Thurman RE, Haugen E, Wang H, et al. (2012): Systematic Localization of Common Disease-Associated Variation in Regulatory DNA. Science 337: 1190–1195.

53. Hatoum AS, Colbert SMC, Johnson EC, Huggett SB, Deak JD, Pathak GA, et al. (2022): Multivariate Genome-Wide Association Meta-Analysis of over 1 Million Subjects Identifies Loci Underlying Multiple Substance Use Disorders. p 2022.01.06.22268753.

54. Lu Z, Gopalan S, Yuan D, Conti DV, Pasaniuc B, Gusev A, Mancuso N (2022, February 11): Multi-ancestry fine-mapping improves precision to identify causal genes in transcriptome-wide association studies. bioRxiv, p 2022.02.10.479993.

55. Donegan JJ, Boley AM, Glenn JP, Carless MA, Lodge DJ (2020): Developmental alterations in the transcriptome of three distinct rodent models of schizophrenia. PLOS ONE 15: e0232200.

