## Supplement for "Multi-omics analyses cannot identify true-positive novel associations from underpowered genome-wide association studies of four brain-related traits"

Indianapolis, IN, USA

8. Department of Biochemistry and Molecular Biology, Indiana University School of Medicine,

Indianapolis, IN, USA

**Contents**

Supplemental Results: pg. 3

Supplemental Table: pg. 4

Supplemental Figures (1-8): pgs. 5-12

**Supplemental Results**

Across both gene prioritization and novel gene discovery analyses, genes identified by more methods in the smaller GWAS tended to have a higher -log10(p) value in the larger GWAS (β=0.22, SE=3.8x10^-3^, *t*=55.6, *p*<2x10^-16^; Supplemental Figure 2), and similarly tended to be identified by more methods in the larger GWAS (β=0.31, SE=3.5x10^-3^, *t*=90.6, *p*<2x10^-16^; Supplemental Figure 3). However, in both cases the effect was larger for gene prioritization than for novel gene discovery (i.e., the interaction between SNP-based significance and number of agreeing methods in the smaller GWAS was significant and positive; y = -log10(p) value in larger GWAS: β=0.35, SE=5.7x10-2, t=6.04, p=1.7x10-9; y = number of significant multi-omics in larger GWAS: β=0.12, SE=1.2x10-3, t=100.2, p<2x10-16).

Follow-up analyses examined every combination of the multi-omics methods (8,178 total combinations), and considered both when combinations of methods agree (e.g., two methods both identify the same gene) and when any of a set of methods agree (e.g., a gene is identified by any of three methods) (Supplemental Figure 4). For novel gene discovery, no combination achieved even a low sensitivity and PPV for any trait (e.g., PPV > 0.2 and sensitivity >0.2). For gene prioritization, no combination of methods consistently out-performed individual methods for all traits. However, in the case of all methods agreeing, the combination of MAGMA and fastBAT out-performed any individual method for SCZ (PPV = 0.69, sensitivity = 0.79). In the case of any of a subset of methods identifying a gene, the combination of fastBAT, MAGMA, H-MAGMA, and FUSION out-performed individual methods for SCZ and AUD/PAU (SCZ: PPV = 0.68, sensitivity = 0.99; AUD/PAU: PPV = 0.92, sensitivity = 0.52).

Follow-up analyses additionally tested whether performance improved if the definition of a ‘significant’ gene in the larger GWAS included those that are proximal to a locus that is trending towards significance (p<0.05x10^-6^) or those that were themselves identified by multi-omics methods in the larger GWAS (p<0.05 bonferroni-corrected). The pattern for gene prioritization did not change when including trending genes in the larger GWAS or when including genes identified by multi-omics in the larger GWAS (Supplemental Figure 5 A&B, E&F). For novel gene discovery, when including trending genes in the larger GWAS, ICV and MDD attained a high PPV with a low sensitivity, when all methods agreed (i.e., the single gene identified by all methods was proximal to a trending locus in the larger GWAS) (Supplemental Figure 5 C&D). Notably, there was no change for AUD/PAU (i.e., the PPV remained nearly 0), and PPV did not exceed 0.5 for SCZ. When including genes identified by multi-omics in the larger GWAS, only MDD attained a high PPV when all methods agreed (i.e., the single gene identified by all methods was also identified by multi-omics in the larger GWAS), while SCZ and ICV did not exceed 0.6, and AUD/PAU remained nearly 0 (Supplemental Figure 5 G&H).

| **Wave** | **Method** | **Trait** | **AUC** | **Specificity** | **PPV** | **Sensitivity** | **NPV** |
| --- | --- | --- | --- | --- | --- | --- | --- |
| **Smaller** | **AdaBoost** |  |  |  |  |  |  |
|  |  | **SCZ** | 0.972 | 0.981 | 0.213 | 0.963 | 0.999 |
|  |  | **MDD** | 0.818 | 0.922 | 0.03 | 0.714 | 0.999 |
|  |  | **AUD/PAU** | 0.78 | 0.989 | 0.02 | 0.571 | 0.999 |
|  |  | **ICV** | 0.95 | 0.99 | 0.6 | 0.9 | 0.999 |
|  | **Average**  **Neural Net** | |  |  |  |  |  |
|  |  | **SCZ** | 0.694 | 0.997 | 0.552 | 0.393 | 0.994 |
|  |  | **MDD** | 0.705 | 0.982 | 0.073 | 0.429 | 0.998 |
|  |  | **AUD/PAU** | 0.783 | 0.994 | 0.04 | 0.571 | 0.999 |
|  |  | **ICV** | 0.649 | 0.998 | 0.15 | 0.3 | 0.999 |
|  | **Ensemble** |  |  |  |  |  |  |
|  |  | **SCZ** | 0.975 | 0.959 | 0.196 | 0.991 | 0.999 |
|  |  | **MDD** | 0.834 | 0.926 | 0.091 | 0.743 | 0.999 |
|  |  | **AUD/PAU** | 0.853 | 0.991 | 0.03 | 0.714 | 0.999 |
|  |  | **ICV** | 0.95 | 0.999 | 0.563 | 0.9 | 0.999 |
| **Larger** | **AdaBoost** |  |  |  |  |  |  |
|  |  | **SCZ** | 0.839 | 0.97 | 0.434 | 0.704 | 0.991 |
|  |  | **MDD** | 0.86 | 0.96 | 0.087 | 0.793 | 0.998 |
|  |  | **AUD/PAU** | 0.75 | 0.99 | 0.136 | 0.509 | 0.999 |
|  |  | **ICV** | 0.857 | 0.999 | 0.666 | 0.714 | 0.999 |
|  | **Average**  **Neural Net** | |  |  |  |  |  |
|  |  | **SCZ** | 0.598 | 0.998 | 0.789 | 0.197 | 0.976 |
|  |  | **MDD** | 0.671 | 0.983 | 0.16 | 0.359 | 0.994 |
|  |  | **AUD/PAU** | 0.696 | 0.996 | 0.21 | 0.396 | 0.998 |
|  |  | **ICV** | 0.606 | 0.998 | 0.15 | 0.214 | 0.999 |
|  | **Ensemble** |  |  |  |  |  |  |
|  |  | **SCZ** | 0.844 | 0.969 | 0.405 | 0.72 | 0.992 |
|  |  | **MDD** | 0.861 | 0.93 | 0.032 | 0.793 | 0.998 |
|  |  | **AUD/PAU** | 0.77 | 0.992 | 0.175 | 0.547 | 0.998 |
|  |  | **ICV** | 0.893 | 0.999 | 0.688 | 0.786 | 0.999 |

**Supplemental Table 1.** Machine learning model performance. Performance of machine learning models in the held-out data from the smaller, earlier, GWAS (Smaller), and in the later, larger GWAS (Larger). AUD/PAU = Alcohol use disorder/Problematic alcohol use; MDD = Major depressive disorder; SCZ = Schizophrenia; ICV = Intracranial volume.

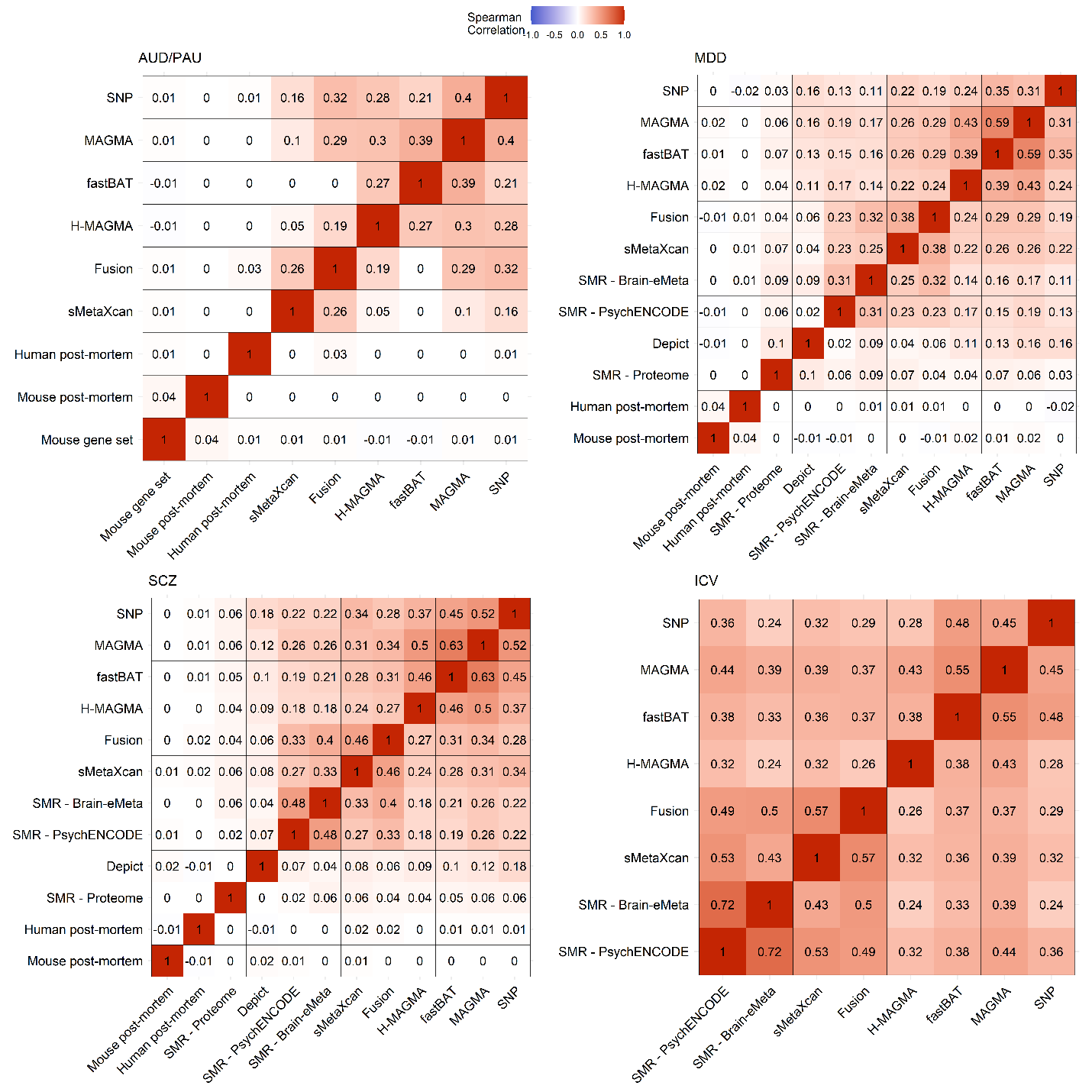

**Supplemental Figure 1**. Spearman correlation between each multi-omics method for each trait. Genes identified by a method are defined as those achieving p<0.05 fdr significance. Methods where no gene passed this threshold are not shown. AUD/PAU = Alcohol use disorder/Problematic alcohol use; MDD = Major depressive disorder; SCZ = Schizophrenia; ICV = Intracranial volume.

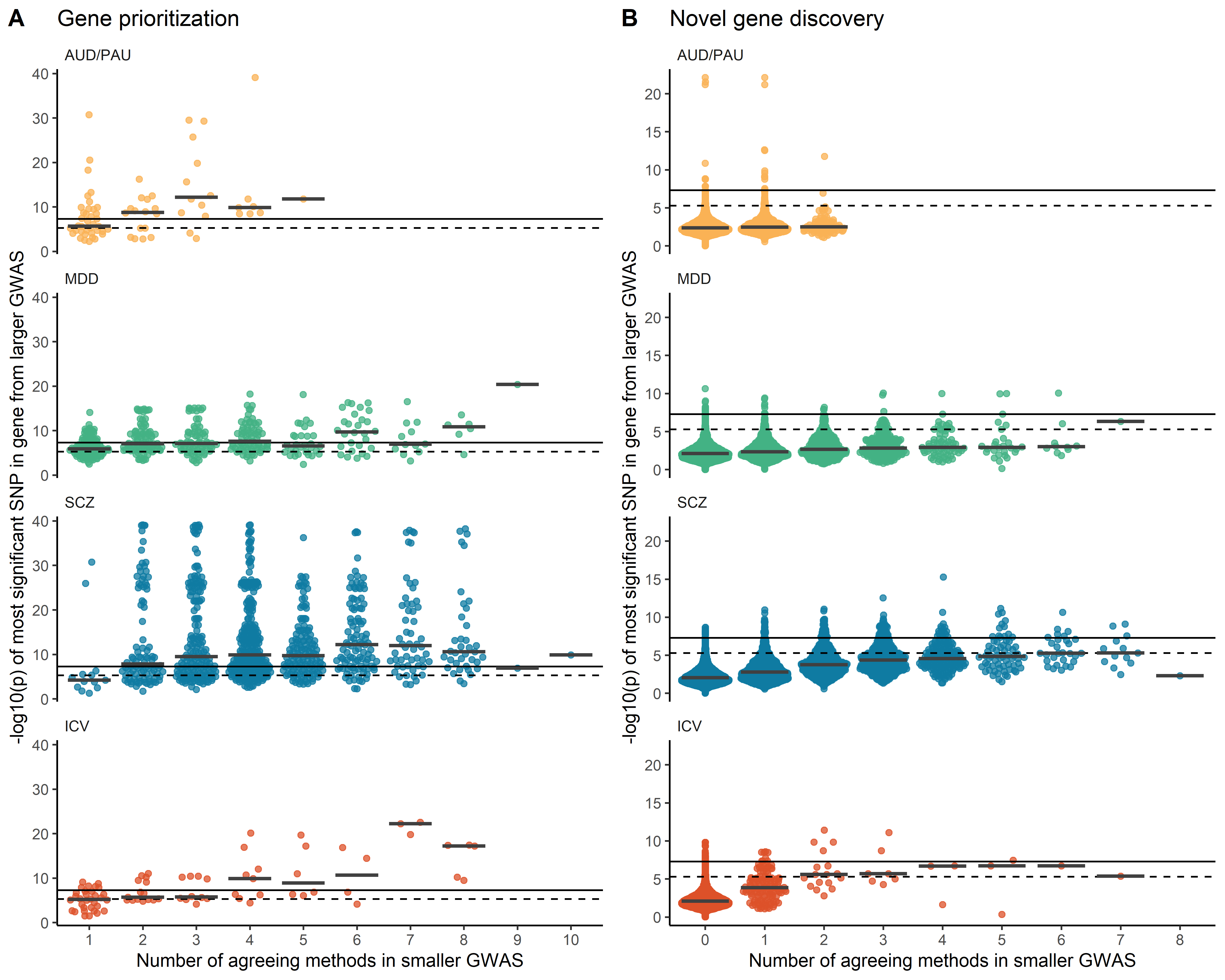

**Supplemental Figure 2**. Prioritized genes have more evidence. Distribution of SNP-level -log10(p)-values for each gene in the larger GWAS, grouped by the number of agreeing methods in the smaller GWAS, for (A) genes that are proximal to a significant locus in the smaller GWAS, and (B) genes that are not.. Solid and dashed lines correspond to p<5x10^-8^ and p<5x10^-6^, respectively. Horizontal bars reflect the median -log10(p) value. AUD/PAU = Alcohol use disorder/Problematic alcohol use; MDD = Major depressive disorder; SCZ = Schizophrenia; ICV = Intracranial volume.

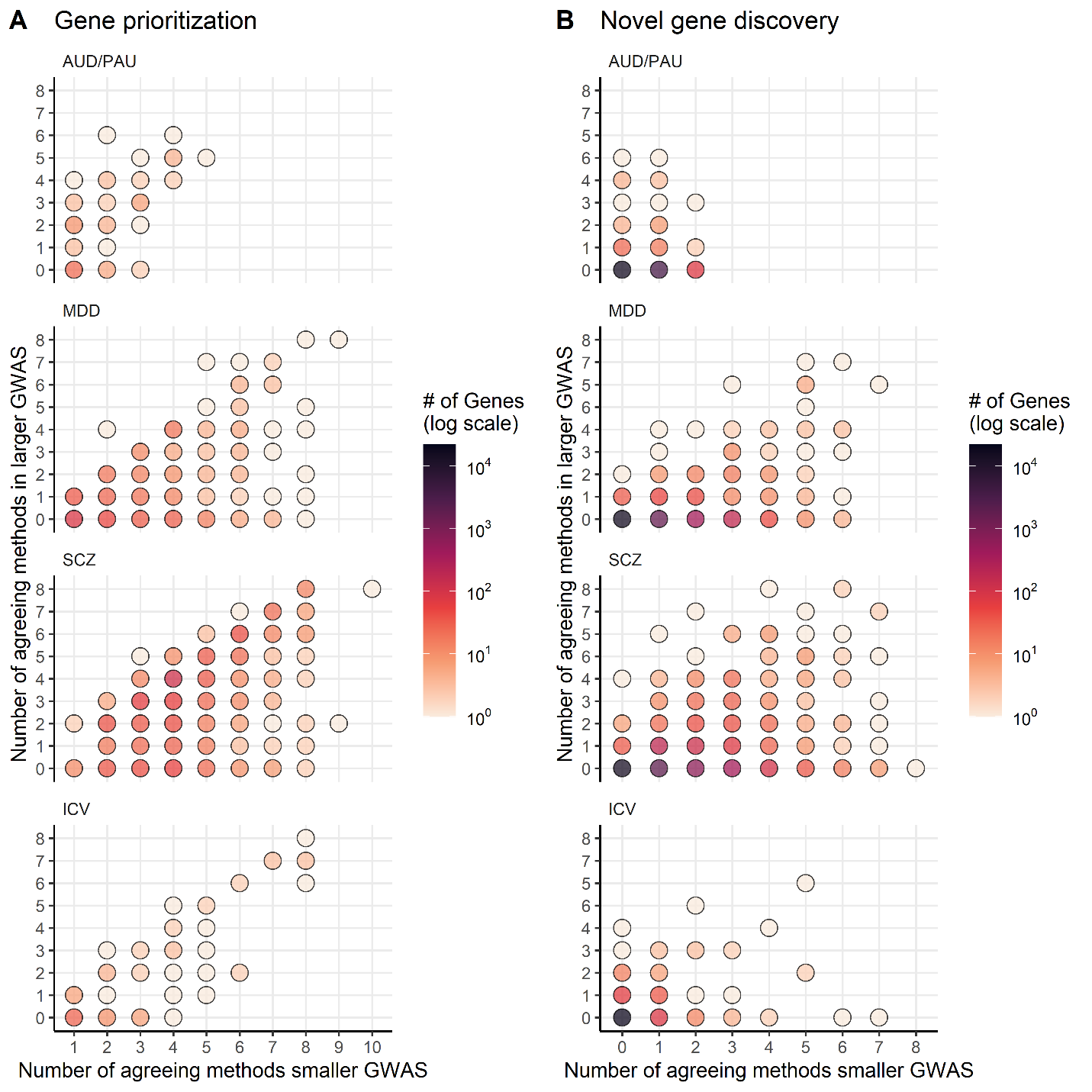

**Supplemental Figure 3**. Genes identified by more multi-omics methods in the smaller GWAS tend to have also been identified by more methods in the larger GWAS. Overlap in genes identified by multi-omics in the smaller and larger GWAS for each trait, grouped by the number of agreeing methods in the smaller GWAS. (A) Genes that are proximal to a significant locus in the smaller GWAS, and (B) Genes that are not. Color reflects the number of genes that lie at the intersection of each level. AUD/PAU = Alcohol use disorder/Problematic alcohol use; MDD = Major depressive disorder; SCZ = Schizophrenia; ICV = Intracranial volume.

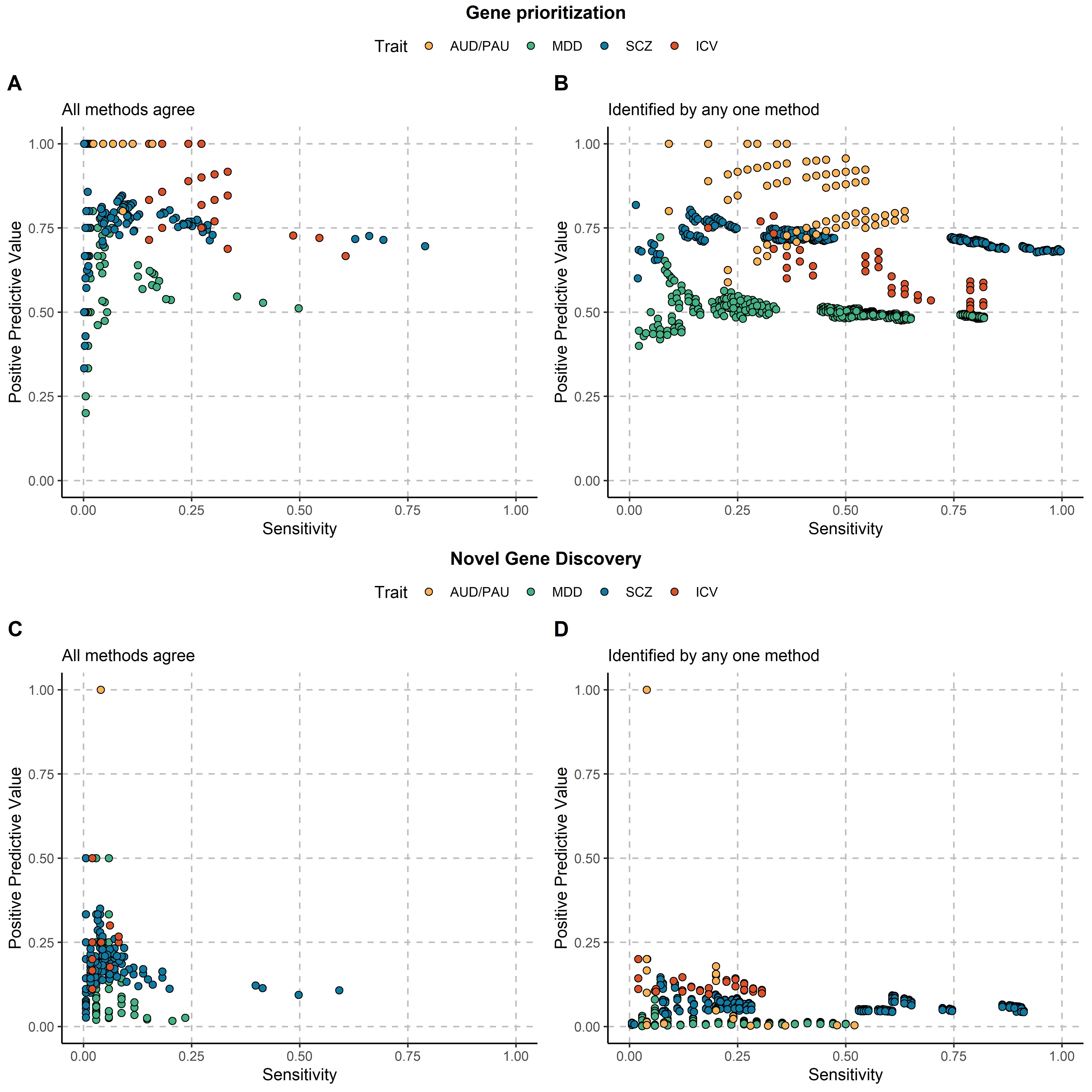

**Supplemental Figure 4**. Performance of every multi-omics method combination. Every possible combination of the 13 methods (from 2 – 13 methods combined) was explored. A) The performance when all methods agree on genes, B) the performance when a gene is tagged by at least 1 of the combined methods. No combination attained both a high positive predictive value and a high sensitivity for any trait (the combination of SNP, MAGMA, and fastBAT performed moderately well only for SCZ). AUD/PAU = Alcohol use disorder/Problematic alcohol use; MDD = Major depressive disorder; SCZ = Schizophrenia; ICV = Intracranial volume.

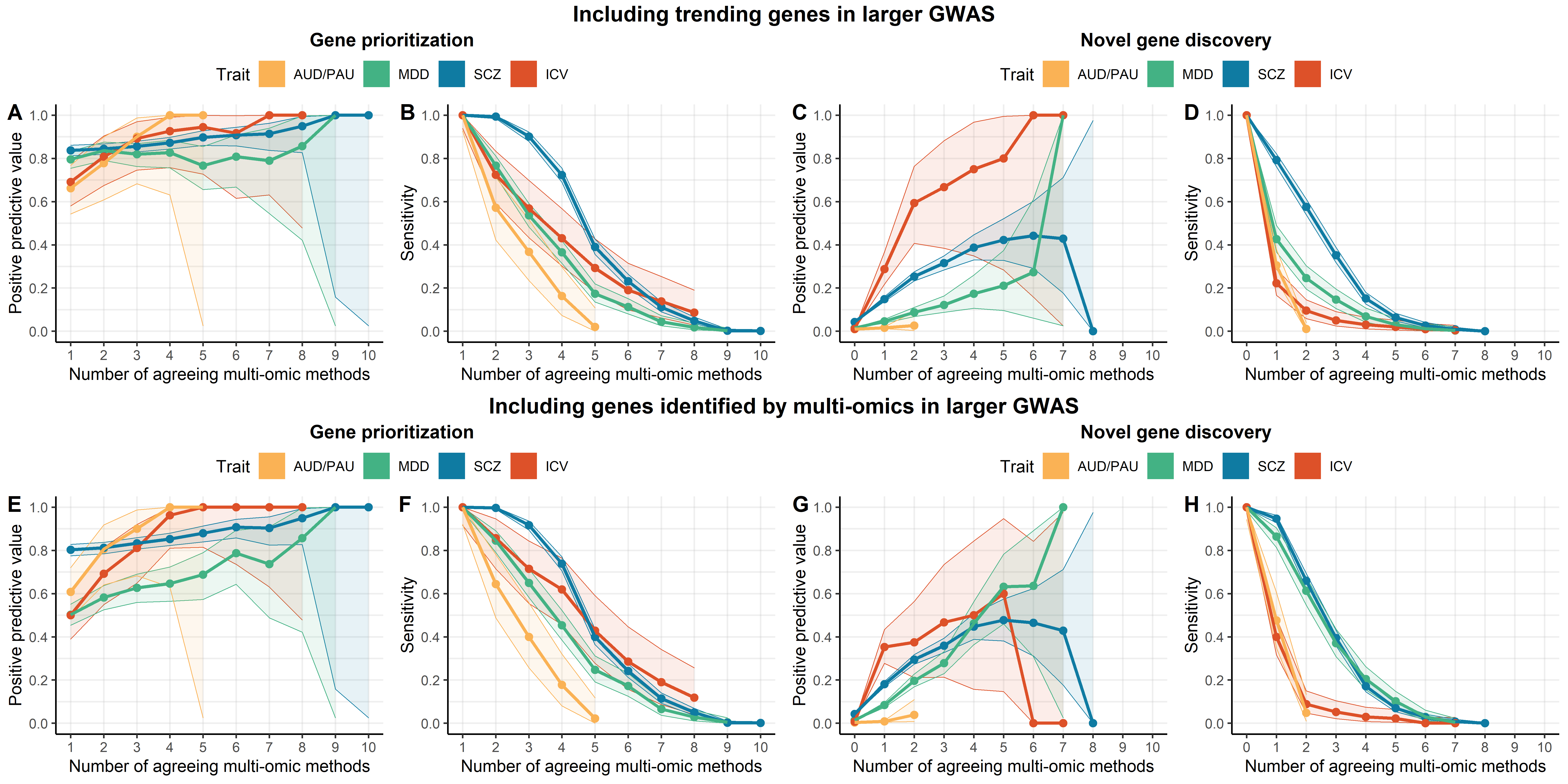

**Supplemental Figure 5**. Positive predictive value and sensitivity when multiple methods agree, under different conditions. A-D: Including trending genes from the larger GWAS (p<5x10^-6^). E-H: Including genes that significant in in any multi-omics methods in the larger GWAS. AUD/PAU = Alcohol use disorder/Problematic alcohol use; MDD = Major depressive disorder; SCZ = Schizophrenia; ICV = Intracranial volume.

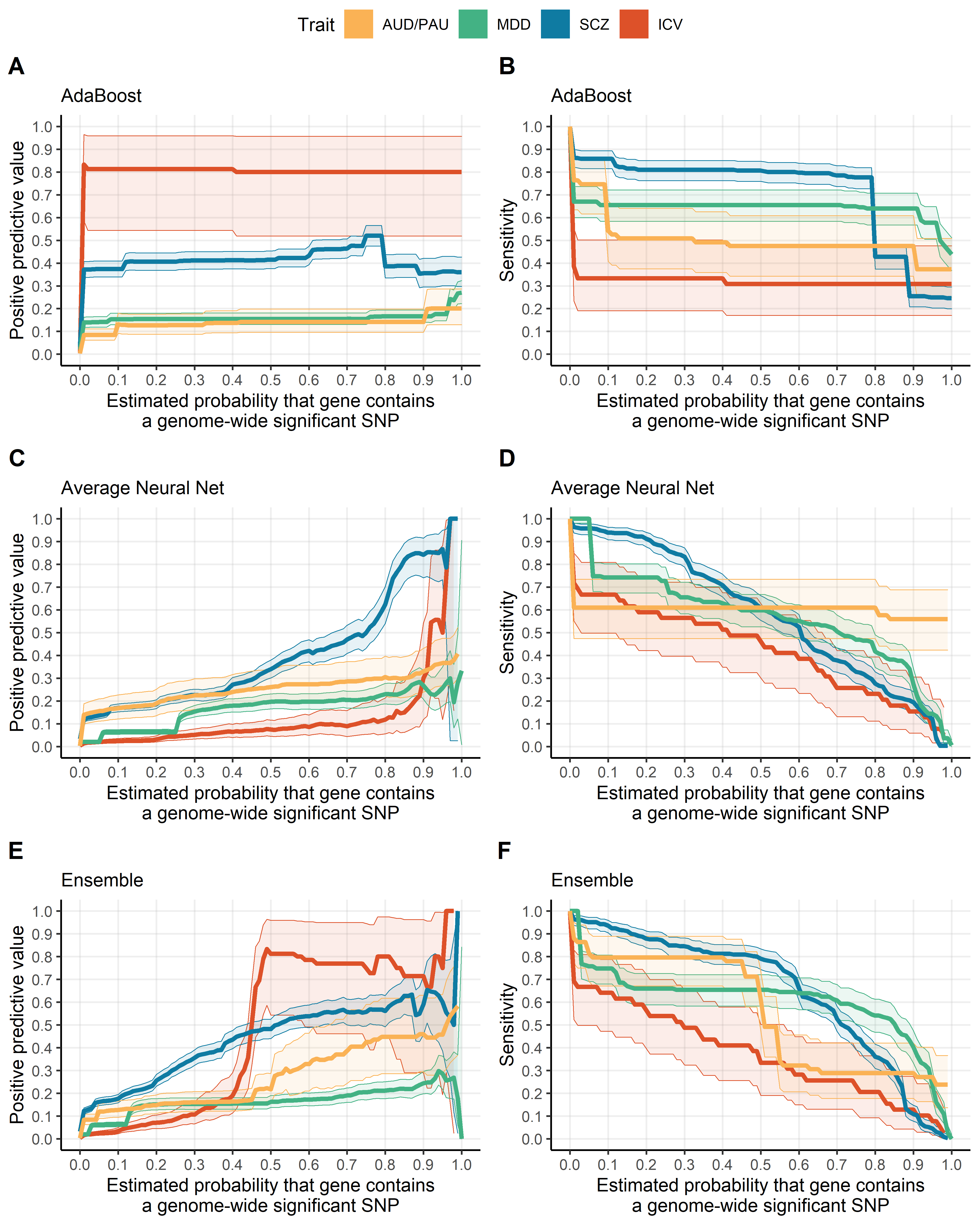

**Supplemental Figure 6.** Machine learning performance in held-out sample from the smaller GWAS. Performance (positive predictive value and sensitivity) of AdaBoost (A&B), Average Neural Net (C&D), and Ensemble (E&F) models in the held-out chromosomes from the smaller GWAS. Models were trained in half the chromosomes in each sample, and tested on the other half. Filled area represents the 95% confidence interval.

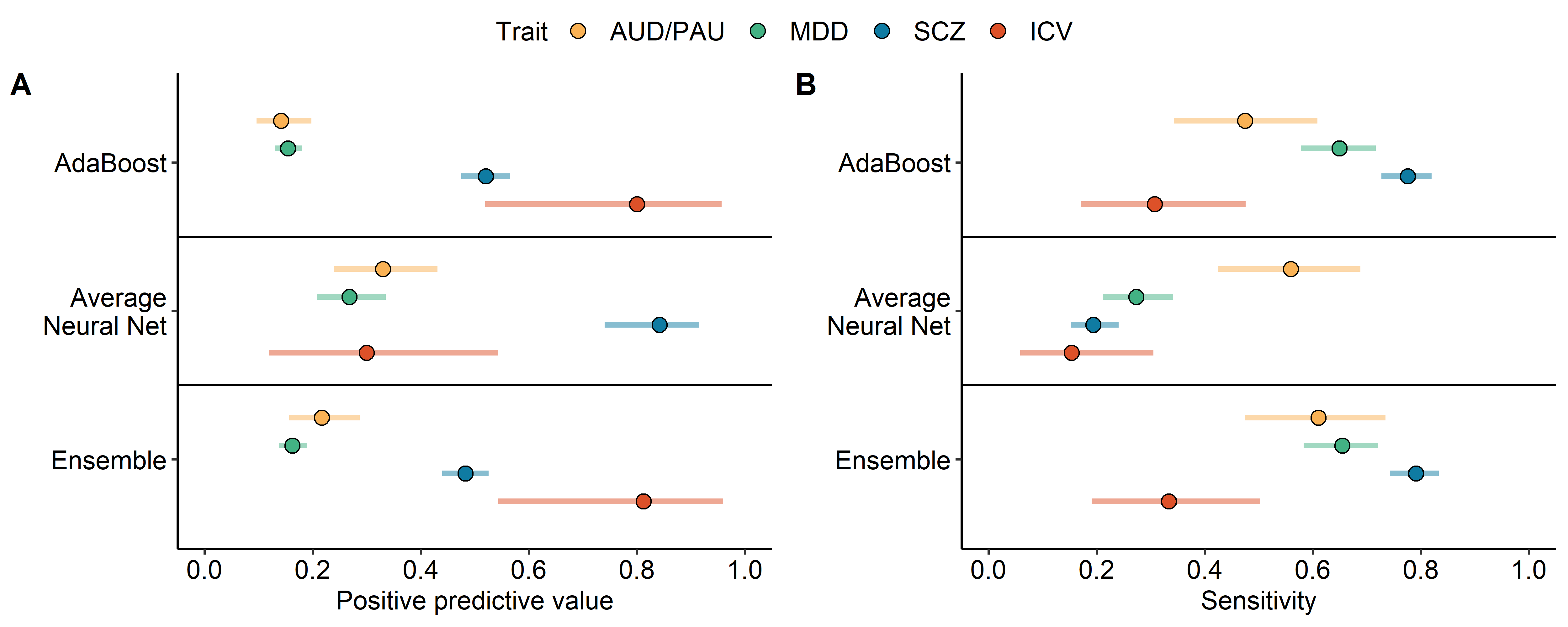

**Supplemental Figure 7.** Machine learning performance in the held-out data from the smaller GWA. Points represent the estimates, while horizonal bars reflect the 95% CI. AUD/PAU = Alcohol use disorder/Problematic alcohol use; MDD = Major depressive disorder; SCZ = Schizophrenia; ICV = Intracranial volume.

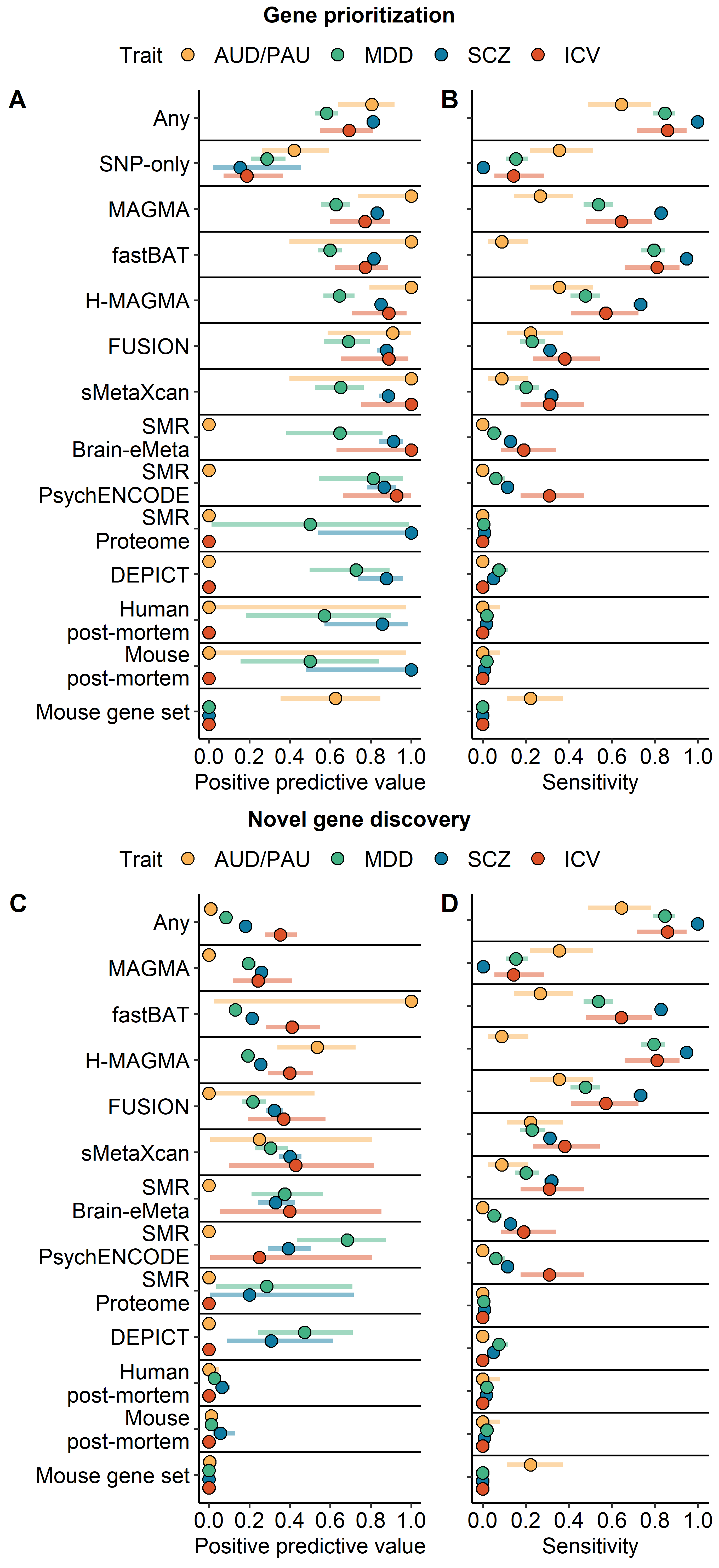

**Supplemental Figure 8**. Performance of multi-omics methods when outcomes include genes identified by multi-omics in the larger GWAS . A&B) The positive predictive value (A) and sensitivity (B) is shown for all methods for all traits. The analysis is restricted to those genes that are proximal to a genome-wide significant locus in the smaller GWAS (i.e., gene prioritization). SNP-only reflects genes that are identified by a significant locus, but not by any multi-omics method. C&D) The positive predictive value (A) and sensitivity (B) is shown for all methods for all traits. The analysis is restricted to those genes that are *not* proximal to a genome-wide significant locus in the smaller GWAS (i.e., novel gene discovery by multi-omics). Points represent the estimates, while horizonal bars reflect the 95% CI. Any = gene is identified by any of the methods. AUD/PAU = Alcohol use disorder/Problematic alcohol use; MDD = Major depressive disorder; SCZ = Schizophrenia; ICV = Intracranial volume.
